# Use treadmills with caution: Walking Energy Expenditure and Metabolic Cost are elevated compared to overground across multiple speeds in healthy young adults

**DOI:** 10.64898/2026.04.14.718406

**Authors:** Sauvik Das Gupta, Kanako Kamishita, Megumi Kondo, Yoshiyuki Kobayashi

## Abstract

**Background:** Treadmill walking is often employed for tightly controlled gait and energetics research, but growing evidence suggests that treadmill-based metabolic and biomechanical measurements may not directly reflect the ecologically valid mode of overground walking. While many previous studies focused on older adults, much less is known about how treadmill walking influences gait energetics and spatiotemporal parameters in young healthy adults across matched speeds.

**Aims and Objectives:** We investigated energy expenditure, metabolic cost of walking and spatiotemporal gait parameters in healthy young adults walking overground and on a treadmill at three speeds (slow-1.0, comfortable-1.3, fast-1.5 m/s). Our hypothesis was that at the comfortable speed, treadmill and overground energetics and gait parameters would be comparable. However, at slow and fast speeds, there would be a significant energetic penalty, accompanied by significant differences in spatiotemporal parameters.

**Methods:** Twenty young participants (10 males and 10 females) completed a randomized cross-over walking protocol with a minimum of ten minutes treadmill familiarization at 1.3 m/s. Breath-by-breath oxygen consumption 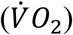 and Respiratory Exchange Ratio were measured using a portable indirect calorimetry system and gait parameters were calculated from Inertial Measurement Units. Gross and net energy expenditures, costs of walking, cadence, average step and stride lengths were calculated. A three-way mixed ANOVA was used for primary statistical analyses.

**Results:** Treadmill walking was characterized by higher gross and net energy expenditures and metabolic costs (p<0.001, η_p_^2^ = 0.6) across all speeds compared to overground. It was also characterized by faster cadence and shorter average step and stride lengths (p<0.001, η_p_^2^ = 0.9). Additionally, there was an effect of sex (p = 0.01, η_p_^2^ = 0.3) on the gait parameters, with females exhibiting a faster cadence and shorter average step and stride lengths than males.

**Discussion and Conclusions:** Our findings show that treadmill walking imposes a medium-to-large metabolic penalty even in healthy young adults, with compensatory gait adaptations, possibly reflecting increased stabilization demands and altered neuromuscular control strategies. These results underscore the limits of generalizing treadmill derived gait data to overground walking and we caution against the uncritical use of treadmills, especially while trying to understand ecologically relevant human walking mechanics and energetics.

## Introduction

Human walking is a fundamental aspect of locomotion that helps us to go from one location to another while minimizing metabolic cost, maintaining stability and optimizing task performance. Walking economy or the Metabolic Cost of Walking (MCoW) is defined as the amount of energy used per unit of distance traveled and generally expressed after body mass normalization. While overground walking is the ecologically valid way of walking, treadmills are extensively used in laboratories as a proxy for daily-life walking. This is because multiple speeds can be tightly controlled and continuous metabolic and biomechanical data can be easily measured. However, it remains ambiguous whether humans, in general, adopt an altered and less economical gait on treadmills and consequently have an elevated MCoW across speeds. Looking at the published literature in this field, we see that a consistent elevation in MCoW for treadmill walking compared to overground is reported in Older Adults (OA). At the same time, there is a lack of published studies involving Young Adults (YA) and the results are inconsistent, compared to those involving OA. Therefore, it is imperative to compare MCoW and gait parameters between overground and treadmill walking across a range of matched speeds in young males and females to better understand the reasons for elevated MCoW in this population.

In general, MCoW is either expressed as the Gross Cost of Walking (GCoW) which reflects the total amount of metabolic energy consumed at the whole-body level or in terms of Net Cost of Walking (NCoW) which is calculated by subtracting the Resting Metabolic Rate (RMR) from the Gross Energy Expenditure (GEE) (Das Gupta et al., 2019). Many past studies (Peterson & Martin, 2010; Hortobagyi et al., 2011; Mian et al., 2006; Pincheira et al., 2017; among others) have consistently shown that OA increase antagonistic muscle co-contraction when walking on a treadmill and this is associated with an elevated MCoW. Parvataneni et al., (2009) reported higher oxygen consumption at matched speeds in OA, whereas Berryman et al., (2012) observed higher MCoW across multiple speeds in OA, on treadmills compared to overground. Furthermore, Martin & Li, (2017) have shown similar elevated treadmill MCoW at same speeds even in YA, though the results in YA are less and more equivocal (Galloway et al., 2021; Daniels et al., 1953; Vickery-Howe et al., 2021; Yngve et al., 2003). In previous work, Das Gupta et al., (2021) found no NCoW difference at same Preferred Walking Speed (PWS) between treadmill and overground in YA, although OA did show an elevation. Very recently a comprehensive systematic review and meta-analysis by Vickery-Howe et al., (2023) showed that treadmill walking in healthy YA and middle-aged adults significantly increases relative oxygen consumption and produces significant differences in kinematics and kinetics with marked increases in cadence while reducing stride and step lengths. Taken together, there are many differences between overground and treadmill walking energetics and so we cannot presume metabolic similarity between the two modes of walking.

From the published literature, we see that beyond metabolic differences, overground and treadmill walking are also biomechanically different, even when the speeds are matched. Treadmill walking is often characterized by higher cadence, shorter step and stride lengths, altered gait variability, and increased demands on balance and sensorimotor control (Alton et al., 1998; Dingwell et al., 2001; Warabi et al., 2005; Riley et al., 2007; McAndrew Young & Dingwell, 2012). Several other studies also explored kinematics, stability constraints and short-term control of gait parameters on treadmills and found that they are different compared to overground (Lee & Hidler, 2008; Reynard & Terrier, 2014; Kozlowska, Latka & West, 2017). All these treadmill specific gait features possibly arise due to differences in visual flow, proprioceptive and vestibular control, compared to overground. All in all, both from a metabolic and biomechanical perspective, there are several differences between overground and treadmill walking and so this shows that the apparent equivalence that is assumed between daily-life overground and treadmill walking is not correct. As a result, it is important to know the true metabolic and biomechanical gait differences between overground and treadmill walking, which warrants further empirical research.

In addition to these commonly known differences, a much-understudied source of variability in treadmill walking studies arise from sex-related differences. Especially due to a shorter height and smaller legs, females typically walk with higher cadence and shorter step and stride lengths than males at matched speeds (Ko et al., 2011; Abualait & Ahsan, 2021; Rayner et al., 2024). Despite these differences, it remains unclear whether males and females respond similarly to the altered biomechanical and neuromuscular control constraints imposed by treadmill walking, compared to overground. Since it is well known that MCoW is more closely related to muscle force production, force-rate, and stabilization demands than to kinematics alone (Kram & Taylor, 1990; Umberger et al., 2003), similar (treadmill-specific) gait adaptations across sexes may nevertheless result in different metabolic consequences. Consequently, it becomes important to look at sex related differences since treadmill-specific constraints could interact with sex-related factors and failing to account for them would obscure actual gait differences between overground and treadmill walking.

Considering the existing literature and importantly to resolve the equivocal differences in YA, we wanted to answer whether young, healthy adults incur a significant metabolic penalty on treadmills compared to overground across a range of speeds, and if so, can we explain it from some basic differences in spatiotemporal parameters? Additionally, we wanted to see if we find significant anthropometrical differences between young males and females, can we also see significant differences in their MCoW and gait parameters? In this study we had a few confirmatory hypotheses and explorative premises. To capture a range of walking speeds, we decided to have three walking speeds and defined them as: Slow - 1.0 m/s, Comfortable - 1.3 m/s and Fast - 1.5 m/s. Our confirmatory hypotheses were as follows:-

- **H1**: Gross and Net Energy Expenditures (GEE and NEE) will increase with increasing walking speeds;
- **H2**: at the comfortable speed, which is close to the PWS of humans, there will be no significant difference in GEE, NEE, MCoW and spatiotemporal gait parameters between treadmill and overground and
- **H3**: at the slow and fast speeds, treadmill walking will show significant elevations in GEE, NEE and MCoW, accompanied by significant differences in spatiotemporal gait parameters.

Exploratively, we posited that if we see significant differences in basic anthropometrics between the young males and females, there would be sex-specific differences in spatiotemporal gait parameters, GEE, NEE and MCoW between treadmill and overground walking.

## Methods

### Participants and Ethics Declaration

10 healthy young Japanese males (mean age 23.3±1.4 years) and 10 healthy young Japanese females (mean age 21.0±1.9 years) were recruited for this study. The sample size was estimated and kept in the range of our previous studies (Das Gupta et al., 2021, 2023) and the other likewise studies cited in Das Gupta et al., (2021). Furthermore, the current study was part of a larger data collection campaign on energy expenditure modeling from wearable Inertial Measurement Units (IMUs) and we followed a recent study (Slade et al., 2021) to estimate our sample size. All participants were free from any specific medications, diabetes, neurological, musculoskeletal and cardiovascular diseases, past surgeries or lower-limb prostheses and had not experienced a fall in the past 6 months. The participants were able to perform regular daily chores without any assistance and were physically active. None of them were undergoing any special strength/endurance or gymnasium training programs. All participants were briefed about the experiments and their participation was completely voluntary and they could leave the experiment at any time. All of them gave written informed consent. Before starting the experiments, basic anthropometrics were measured – body mass, height, waist circumference, dominant hand and leg, and Lower Limb Lengths (LLL) from the greater trochanter to malleolus and foot. Body Mass Index (BMI) and Waist-to-Height ratio were calculated from the measured variables. The ergonomics ethical review committee of the National Institute of Advanced Industrial Science and Technology (AIST, Japan) approved the experimental protocol (Reference number: 2024-1431, Approval number: 73002030-E-20240617-001) in accordance with the principles stated in the Declaration of Helsinki.

### Experimental Design & Protocol

#### Pre-experiment factors

Before any measurements were taken, we advised the participants to have only a light breakfast or lunch (moderate intake of fats and carbohydrates and minimal intake of protein and fiber-rich foods) and to not consume any alcohol or caffeine-related products.

They were specifically told not to smoke tobacco until the end of the experiments. We also ensured that at least 2 hours of time was elapsed from their last food intake to any metabolic data measurement, to minimize the thermic effects of food. For the overground walking trials, the participants walked along an oval track, with 5 m straights, taking smooth turns at the end of the tracks, inside a laboratory (see: Figure 1). For the treadmill walking trials, a floor-level treadmill (T101, Horizon Fitness Inc., USA) was used which was placed at approximately 20 cm from the ground (wheels to belt). Handrails were not provided during the treadmill walks and free arm swing was ensured. To minimize fatigue, we gave at least 4 minutes of seated rest between every walking trial and at least 10 minutes of seated rest while changing from one walking mode to another. We pseudo-randomized the walking speeds and half of the participants completed the overground walking trials first and the other half completed the treadmill walking trials first (see: Figure 1).

**Figure 1:**
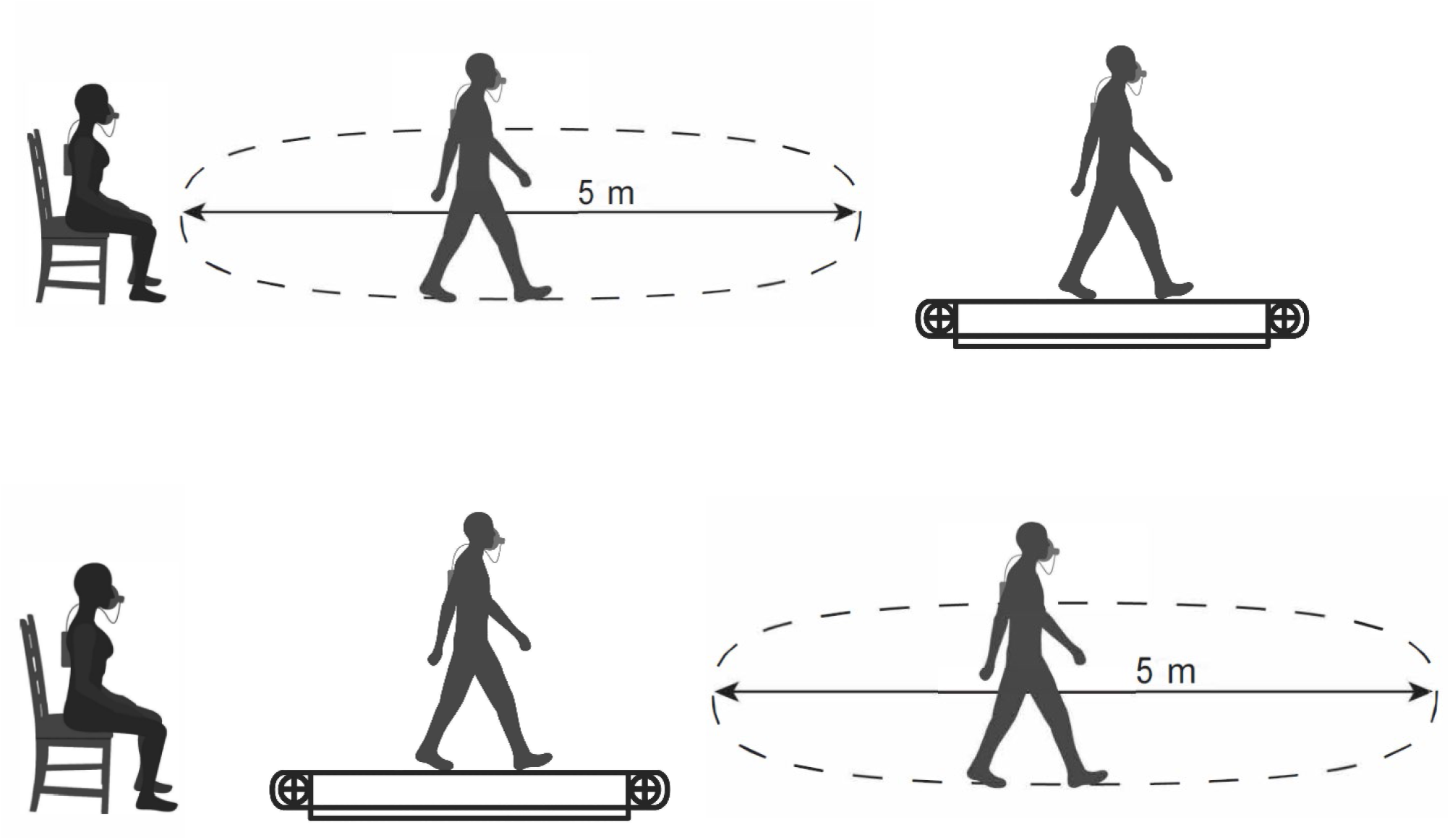
Schematic overview of the energy expenditure measurements. Resting Metabolic Rate was measured while the participants were seated for 5 min on a chair, at the beginning of the experimental session. Metabolic Cost of Walking was measured which included an overground trial and a treadmill trial, for all the three walking speeds. During the overground trial, participants walked continuously along the schematically drawn oval track with 5 m straights and taking smooth turns at the end of the track. During the treadmill trial, the target speeds were set and the participants walked on the treadmill at those speeds. Half of the participants (N=10) completed the overground trial first and then the treadmill and the other half (N=10) did vice-versa.

#### Preparation

The participants were fitted with 4 IMUs (Movella DOT, Movella Inc., USA) on the right thigh, right shank, left thigh and left shank and the data were collected at a sampling rate of 60 Hz. This sampling rate was chosen to enable real-time streaming of the IMU data to the dedicated mobile application, to be checked for any issues by one of the experimenters. A schematic diagram is shown in Figure 2. However, in this study we analyzed only the right thigh and right shank IMU data as the right side was the dominant side for all the participants. They were also fitted with a facemask (Hans Rudolph Inc., USA) and a portable Mobile Aeromonitor AE-100i (Minato Medical Science Co., Ltd., Japan) to measure rates of oxygen consumption and carbon dioxide liberation. All the devices were properly calibrated every day, before the start of each experiment, according to the manufacturer’s guidelines. All the participants were provided with Mizuno Wave Rider shoes (Mizuno Corporation, Japan), according to their foot sizes, by the research team at AIST.

**Figure 2:**
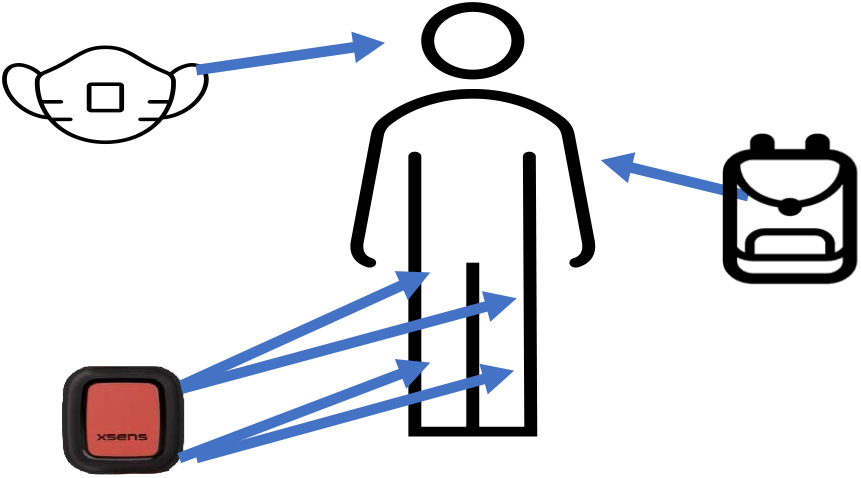
Schematic overview of the IMU sensor and the wearable indirect calorimetry system. 4 IMUs were attached on the right thigh, right shank, left thigh and the left shank of each of the participants. For the metabolic measurement, the participants carried the metabolic device as a wearable backpack and wore facemasks. It was ensured that there was no leakage of air (inspired/expired) from the facemasks.

#### Data collection

Though there were some participants who had previous treadmill running experience, none of them had prior treadmill walking experience. Nevertheless, we decided to familiarize each of our participants for at least 10 minutes on the treadmill at the Comfortable Walking Speed (CWS) of 1.3 m/s in the beginning. After that we gave them 5 minutes of seated rest and then they were fitted with the IMUs, the facemask and the mobile aeromonitor. Before the start of any of the walking trials, we measured the seated RMR (see Figure 1) for 5 minutes (see Das Gupta et al., 2021, 2023). After the resting measurement, all of them walked for at least 6 minutes at each of the walking speeds on both overground and treadmill. For the overground trials an experimenter timed the participants’ walking time of the track with a stopwatch during their walks in the 5 m overground track and ensured that within the first 2 minutes of the walks, all the participants were walking at the desired speed condition. If needed, verbal encouragements were given in native Japanese to ensure that the participants kept walking at the target speeds. In this way we ensured that after they had achieved steady state and during the last 4 minutes of every walking trial, the participants were all walking at the desired speeds. The metabolic steady state was further visually confirmed by checking the wirelessly transmitted data from the mobile aeromonitor to its dedicated laptop, during the resting and the walking trials. At the end of the overground trials, the participants were verbally told to stop walking and then all the measurement devices were stopped. For the treadmill trials we increased the speed from 0 m/s to the desired speed within the first 1 minute of the start of the walk and at the end, the treadmill speed was slowly reduced from the set speed to 0 m/s again within 1 minute and then all the measurement devices were stopped.

#### Calculation of the Energy Expenditure and Metabolic Cost of Walking and related gait parameters

For the resting trial, we rejected the first and last 30 seconds of data from the 5 minutes of constant sitting and calculated the RMR. The RMR (in J kg^−1^ s^−1^ or Watts kg^−1^) was calculated using the resting Respiratory Exchange Ratio (RER, unitless) and the rate of resting oxygen consumption 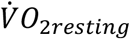, in l of O_2_ kg^−1^ s^−1^) using the Lusk equation (Lusk 1923). For every walking trial, we did not analyze the first 3 minutes of data to ensure a metabolic steady state and the last 30 seconds of data as it was used to stop all the measurement devices. From the steady state region of the walking trials, all the necessary metabolic variables were calculated after averaging. The Gross Energy Expenditure (GEE, in J kg^−1^ s^−1^ or Watts kg^−1^) was calculated using the walking RER (unitless) and the rate of walking oxygen consumption (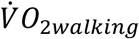, in l of O_2_ kg^−1^ s^−1^) using the same Lusk equation. From the GEE we subtracted the RMR (in J kg^−1^ s^−1^ or Watts kg^−1^) to get the Net Energy Expenditure (NEE, in J kg^−1^ s^−1^ or Watts kg^−1^). Then we divided the GEE and NEE by the Walking Speed (WS, in m s^−1^) to get the Gross Cost of Walking (GCoW) and the Net Cost of Walking (NCoW) in the units of J kg^−1^ m^−1^. These were the primary variables of interest in this study.

Additionally, we calculated a few spatiotemporal parameters. The total number of gait cycles of every walking trial was calculated from the shank IMU data (from the vertically oriented z-axis, sagittal plane angular velocity from the gyroscope) using an open-source algorithm from Slade et al., (2021) assuming steady-state bilateral symmetry. We used the algorithm for treadmill walking as in Slade et al., (2021) and extended it for overground walking in this study. Although (in-lab) overground walking generally involves taking turns in addition to straight-line walks, it has been previously reported (Romijnders et al., 2021) that shank IMUs are also valid for detecting gait events during turns in overground walking, with high accuracy. Furthermore, as we knew the total walking times for each completed walking trial, we could directly calculate the Cadence (steps s^−1^). After that we used the Cadence and the walking speed (in m s^−1^) to calculate the average step length (m) and then doubled it to calculate the average stride length (m). All the equations used are described as follows:-

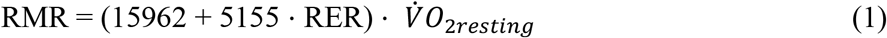

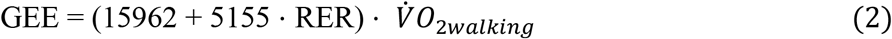

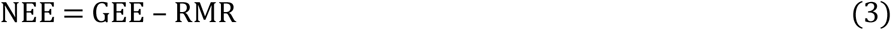

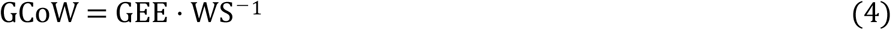

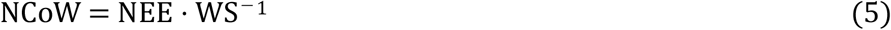

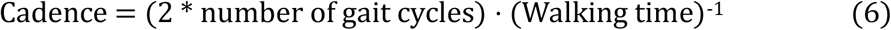

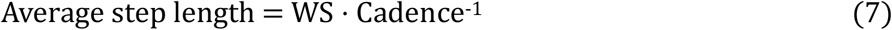

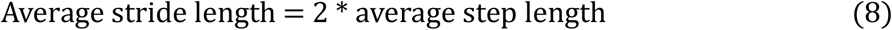

#### Statistical tests

For age, anthropometric parameters, RMR, GEE, NEE, GCoW, NCoW, Cadence, average step and stride lengths, first we checked for normality using the Shapiro-Wilk test. For the RMR, age and anthropometric parameters we conducted independent samples (between-groups) T test (Student’s or Mann-Whitney) between the males and females in our cohort. We also performed Levene’s test to check for the equality of variances. The effect sizes for the statistically significant effects were reported as either Hedge’s G or Rank Biserial correlation.

Since in this study we wanted to look at and explain the differences in GEE, NEE, GCoW and NCoW between overground and treadmill with differences in the spatiotemporal parameters, we mainly chose to look at the intra-individual (within-group) differences. We then conducted a three-way mixed ANOVA with sex (males versus females) as the between-groups factor, walking surface (overground versus treadmills) and walking speeds (slow, comfortable, fast) as the within-groups factor. Mauchly’s test was used to do sphericity corrections and Greenhouse-Geisser corrections were applied, when necessary. We performed Levene’s test to further check for the equality of variances for the remaining variables. If there were statistically significant main and interaction effects, only for those effects, we further conducted post-hoc tests with Bonferroni’s corrections. The effect sizes for the statistically significant effects are reported as partial eta-squared (η_p_^2^). The open-source statistical software JASP (version 0.18.3.0) was used for all the statistical tests and the default value of α=0.05 was chosen as the level of statistical significance.

## Results

### Age, Resting Metabolic Rate and Anthropometric measurements

The results of the RMR, age and anthropometric parameters are given in Table 1, where the mean, standard deviations (mean ± SD) and the effect sizes are mentioned. The effect sizes are calculated as the male group, subtracted from the female group. The statistically significant differences (p<0.05) between the males and females are denoted with an asterisk (*) and the corresponding p values are given. The young Japanese males were heavier, taller, with a larger waist circumference and longer lower limb lengths. The females were also significantly younger than the males, but the mean age values indicate that all the participants were young.

**Table 1:** Age, RMR and anthropometric parameters of the study participants.

| **Variables** | **Females**<br>**(N=10)** | **Males**<br>**(N=10)** | **Effect Size** | **Statistically**<br>**significant**<br>**(with p values)** |
| --- | --- | --- | --- | --- |
| RMR (J kg<sup>-1</sup> s<sup>-1</sup>) | 1.30 ± 0.13 | 1.36 ± 0.10 |  |  |
| Age (years) | 21.00 ± 1.94 | 23.30 ± 1.42 | -0.7 | p=0.01* |
| Body mass (kg) | 49.58 ± 5.39 | 63.20 ± 9.06 | -1.8 | p<0.001* |
| Height (m) | 1.58 ± 0.04 | 1.73 ± 0.05 | -3.3 | p<0.001* |
| Waist circumference (m) | 0.72 ± 0.05 | 0.80 ± 0.07 | -1.12 | p=0.02* |
| LLL up to Malleolus (m) | 0.73 ± 0.03 | 0.80 ± 0.04 | -1 | p<0.001* |
| LLL up to Foot (m) | 0.79 ± 0.03 | 0.86 ± 0.04 | -1 | p<0.001* |
| BMI (kg m<sup>-2</sup>) | 19.91 ± 1.93 | 21.1 ± 2.72 |  |  |
| Waist to Height Ratio | 0.46 ± 0.03 | 0.46 ± 0.04 |  |  |

### Main statistical analyses

#### Gross and Net Energy Expenditures and Metabolic Costs of walking

From the mixed-ANOVA, there were statistically significant main effects ((F(1.4, 25.4) = 70.7 for GEE) and (F(1.4, 25.2) = 78.0 for NEE), p<0.001, η_p_^2^ = 0.8) for walking speeds and walking surface ((F(1, 18) = 23.5 for GEE) and (F(1,18) = 25.4 for NEE), p<0.001, η_p_^2^ = 0.6) for both gross and net energy expenditures. Post hoc tests revealed significant effects (Bonferroni adjusted) between all the three pairs of speeds (slow-comfortable, slow-fast and comfortable-fast, p<0.001) and between overground and treadmill (p<0.001), with the faster speeds and treadmill showing elevated energy expenditures. Due to several statistically significant anthropometric differences between our participants, we are showing the males and females with separate sub-figures in this section representing their GEE, NEE, GCoW, NCoW and the various gait parameters. Figure 3 shows the descriptive plots for males (A,C) and females (B,D) for GEE and NEE.

**Figure 3:**
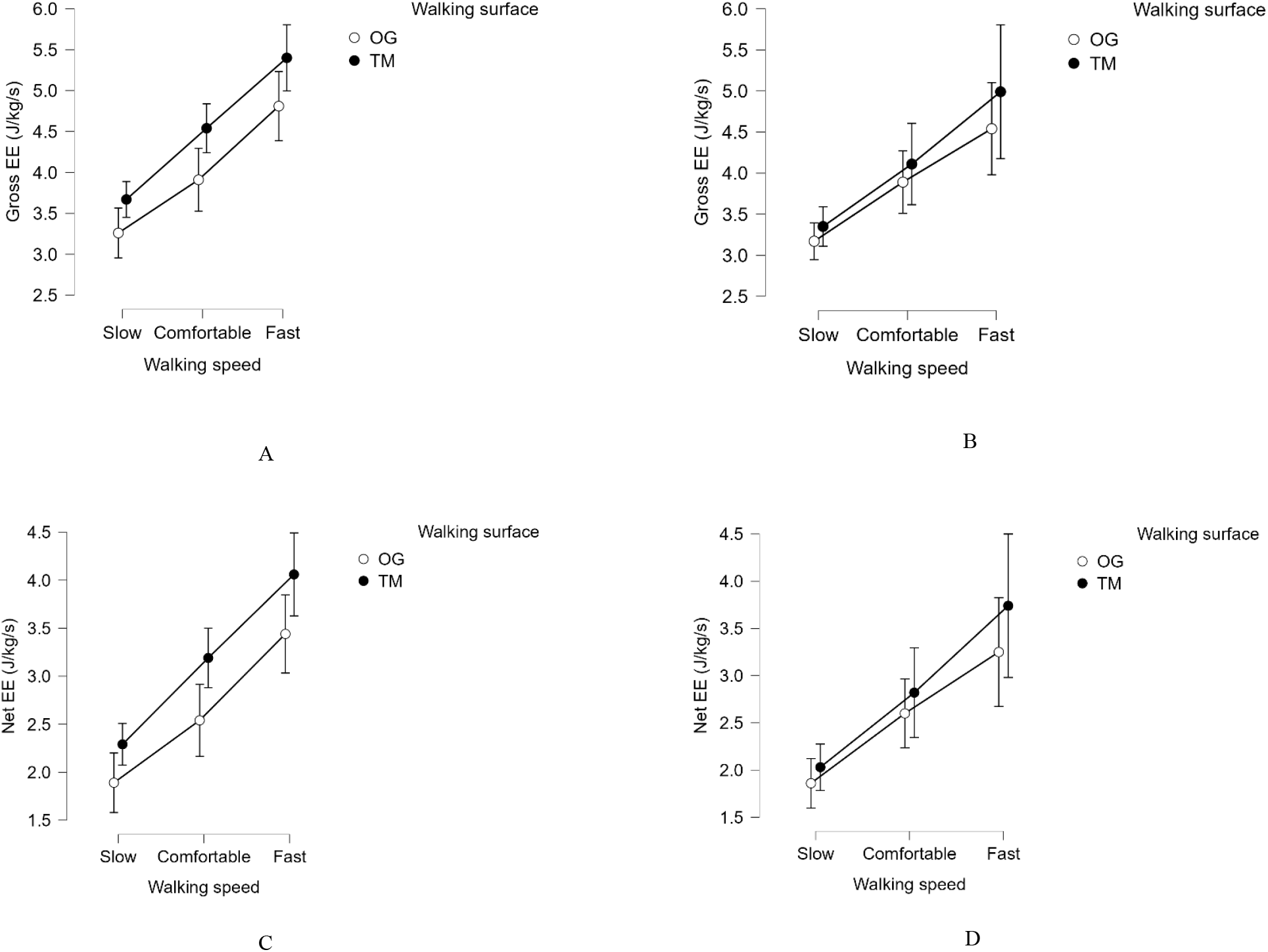
Line-plots of Gross Energy Expenditure (GEE) and Net Energy Expenditure (NEE) are shown for overground (OG, in open white circles) and treadmill (TM, in black closed circles). The circles denote the mean values and the vertical straight black lines across the circles are the error bars denoting the 95% Confidence Intervals (CIs). The x-axis shows the walking speeds in text and the y-axis shows the main metabolic variable and its unit. The top row (A,B) depicts the GEE and the bottom row (C,D) depicts the NEE. The left column (A,C) depicts the data for the young males and the right column (B,D) depicts the data for the young females. It is clear from these figures that at matched speeds, the treadmill incurs a metabolic penalty for both males and females (more for males) and with increasing speeds, expectedly there is an increase in both GEE and NEE.

For gross metabolic costs, there was a statistically significant main effect for the walking surface (F(1,18) = 25.3, p<0.001, η_p_^2^ = 0.6), with post hoc tests indicating that treadmills incurred a greater metabolic cost than overground, for all the speeds (p<0.001). For net metabolic cost of walking, there were statistically significant main effects for walking speed (F(1.5, 27.2) = 9.6, p = 0.002, η_p_^2^ = 0.4) and walking surface (F(1,18) = 24.3, p<0.001, η_p_^2^ = 0.6). Post hoc tests revealed a significant elevation (Bonferroni adjusted) in cost between fast and slow (p<0.001) and fast and comfortable speeds (p = 0.02) and for treadmill compared to overground (p<0.001). Figure 4 shows the descriptive plots for males (A,C) and females (B,D) for GCoW and NCoW.

**Figure 4:**
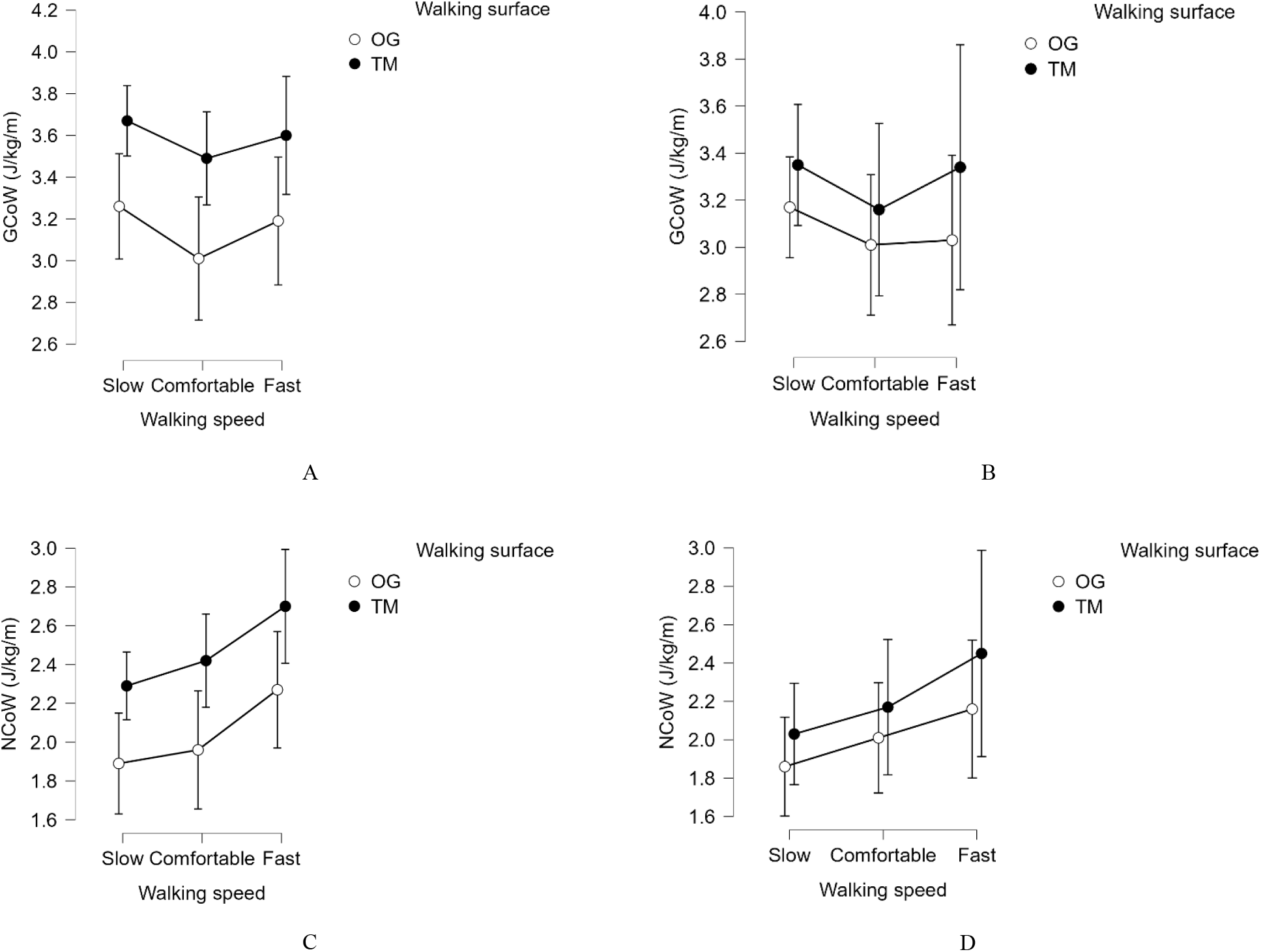
Line-plots of Gross Cost of Walking (GCoW) and Net Cost of Walking (NCoW) are shown for overground (OG, in open white circles) and treadmill (TM, in black closed circles). The circles denote the mean values and the vertical straight black lines across the circles are the error bars denoting the 95% CIs. The x-axis shows the walking speeds in text and the y-axis shows the main metabolic variable and its unit. The top row (A,B) depicts the GCoW and the bottom row (C,D) depicts the NCoW. The left column (A,C) depicts the data for the young males and the right column (B,D) depicts the data for the young females. It is clear from these figures that at matched speeds, the treadmill incurs a metabolic penalty for both males and females (more for males). With increasing speeds, the ecologically valid GCoW shows the classic and expected U-shaped relationship with speed, with an energetic minimum near the CWS. On the contrary, due to the same baseline subtraction of the RMR from the GEE, to calculate the NEE and hence NCoW, we see a flattening of NCoW for the slow and comfortable speeds and an increasing NCoW towards the fast speed

#### Spatiotemporal Gait Parameters

For cadence, there were statistically significant main effects for walking speed (F(1.3, 24.0) = 344.7, p<0.001, η_p_^2^ = 0.95), walking surface (F(1,18) = 122.5, p<0.001, η_p_^2^ = 0.9) and an interaction effect for walking speed*walking surface (F(2,36) = 28.4, p<0.001, η_p_^2^ = 0.6). Post hoc tests revealed a significant difference (Bonferroni adjusted) in cadence between all the speed combinations (p<0.001) and a significant elevation on treadmill compared to overground (p<0.001). For the interaction effect, all the post hoc combinations were statistically significant (p<0.001), with the only exception being the difference between comfortable speed on overground and slow speed on treadmill (p = 0.8), where the mean difference was around 0.05 (95% CI = −0.03 – 0.12) steps/s. The key result was that at matched speeds, all our participants had a faster cadence on the treadmill compared to overground. Finally, there was a statistically significant main effect of sex (F(1, 18) = 7.4, p = 0.01, η_p_^2^ = 0.3), with the post hoc test showing a similar level of significance (p = 0.01). The mean difference revealed that the females walked with a faster cadence than the males. Figure 5 shows the descriptive plots for cadence for males (A) and females (B).

**Figure 5:**
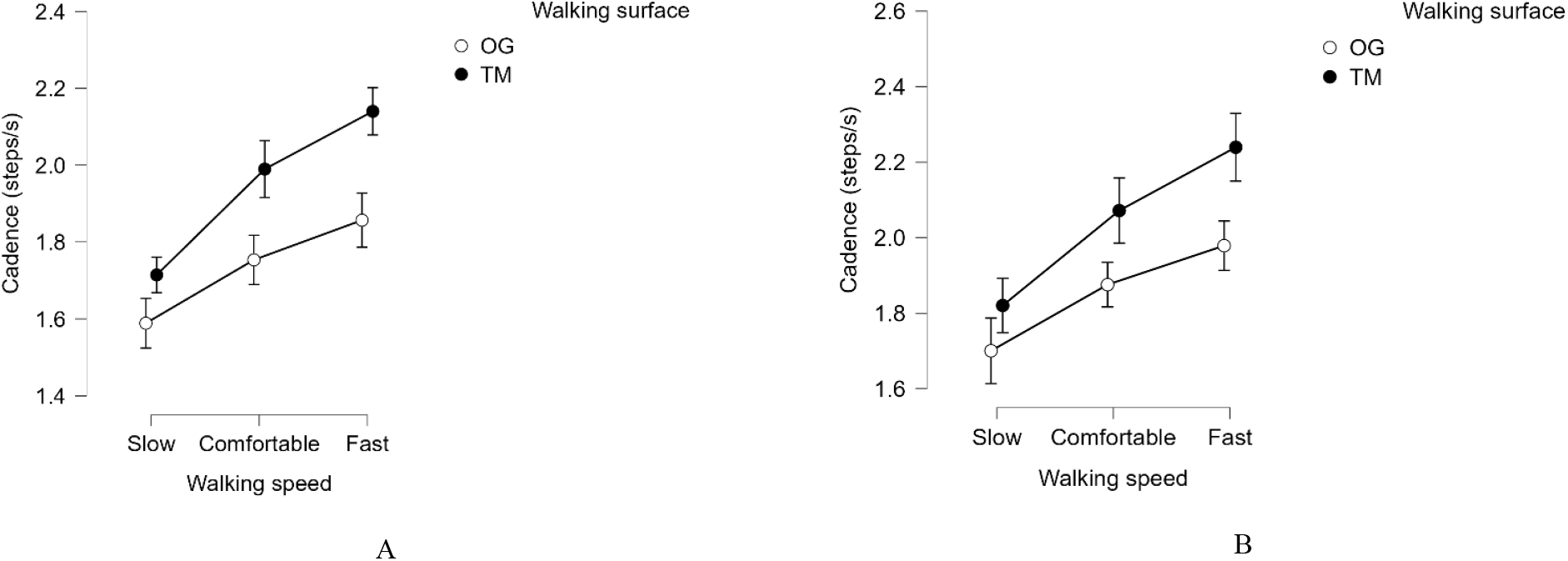
Line-plots of cadence are shown for overground (OG, in open white circles) and treadmill (TM, in black closed circles). The circles denote the mean values and the vertical straight black lines across the circles are the error bars denoting the 95% CIs. The x-axis shows the walking speeds in text and the y-axis shows the gait parameter and its unit. Figure 5A depicts the data for the young males and Figure 5B depicts the data for the young females. It is clear from these figures that at matched speeds, the treadmill forces an increased cadence (quicker steps) for both males and females and with increasing speeds, expectedly there is an increase in cadence. If we compare the y-axis values of figures 5A and B, we can see that the females had a faster cadence than males.

For average step and stride lengths, there were statistically significant main effects for walking speed (F(1.4, 24.4) = 560.0, p<0.001, η_p_^2^ = 0.97), walking surface (F(1,18) = 122.5, p<0.001, η_p_^2^ = 0.9) and an interaction effect for walking speed*walking surface (F(2,36) = 31.2, p<0.001, η_p_^2^ = 0.6). Post hoc tests revealed a significant difference (Bonferroni adjusted) in average step and stride lengths between all the speed combinations (p<0.001) and a significant reduction on treadmill compared to overground (p<0.001). For the interaction effect, all the post hoc combinations were statistically significant (p<0.01). The key result was that at matched speeds, all our participants had smaller average step and stride lengths on the treadmill compared to overground. Finally, there was a statistically significant main effect of sex (F(1, 18) = 7.5, p = 0.01, η_p_^2^ = 0.3), with the post hoc test showing a similar significance (p = 0.01). The mean differences showed that the average step and stride lengths of females were smaller than males. Figure 6 shows the descriptive plots for average step length for males (A) and females (B). Since in this study, the average stride length is mathematically just double the average step length, the figure looks like that of average step length and so is shown in the supplementary information.

**Figure 6:**
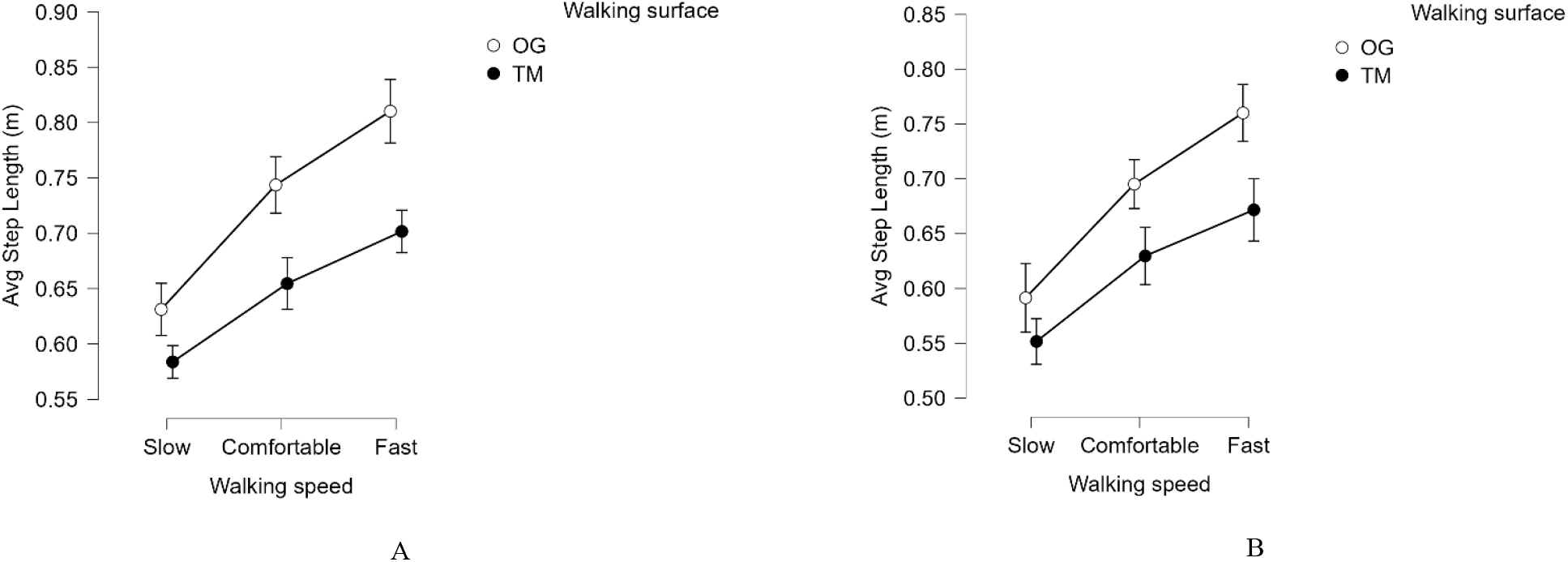
Line-plots of average step length are shown for overground (OG, in open white circles) and treadmill (TM, in black closed circles). The circles denote the mean values and the vertical straight black lines across the circles are the error bars denoting the 95% CIs. The x-axis shows the walking speeds in text and the y-axis shows the gait parameter and its unit. Figure 6A depicts the data for the young males and Figure 6B depicts the data for the young females. It is clear from these figures that at matched speeds, the treadmill forces a reduced average step length (smaller steps) for both males and females and with increasing speeds, expectedly there is an increase in average step length. If we compare the y-axis values of figures 6A and B, we can see that the females had a shorter average step length than males.

## Discussion

Since treadmills are widely used in gait laboratories as a valid proxy for natural, overground walking, in this study we wanted to confirm if healthy YA incur a metabolic penalty on the treadmill compared to overground at matched speeds and if it is accompanied by an altered gait. Additionally, we also wanted to explore if sex differences influence the metabolic and gait parameters on the treadmill, compared to overground. To answer these questions we recruited healthy, young Japanese males and females and measured their energy expenditure, metabolic cost and spatiotemporal gait parameters at three speeds (Slow – 1.0 m/s, Comfortable – 1.3 m/s and Fast – 1.5 m/s) during walking on both overground and treadmill. Our results indicate that our first hypothesis (**H1**) was fully accepted. Contrary to our expectations, the second hypothesis (**H2**) was rejected as even at the CWS the treadmill incurred an energetic penalty, marked with an altered gait. Our third hypothesis (**H3**) was also fully accepted. Additionally, our explorative suggestion was partially confirmed. Our data clearly demonstrate that treadmill walking, even in YA, imposes a metabolic penalty and elicits an altered gait relative to overground walking across multiple speeds, which is less economical and thus, indicative of a sub-optimal strategy. Below we will discuss our results in detail and elucidate them by considering the previous work in the literature.

The key finding of this study was that GEE, NEE and NCoW increased with walking speed and were higher on the treadmill, whereas GCoW was higher on the treadmill and exhibited a U-shaped relationship with speed. This is expected as walking faster requires greater mechanical power and muscle activations, while treadmill walking imposes additional constraints of enclosed space, different neuromuscular and stabilization requirements and sensory-motor mismatch than overground walking (Martin & Li, 2017). This results in elevated energy expenditures and metabolic cost, even at matched speeds (Kram & Taylor, 1990; Donelan et al., 2002).

Accompanied by these metabolic penalties, for both overground and treadmill walking, we saw significant elevations in cadence and average step and stride lengths with increasing walking speeds. However, at matched speeds, treadmill walking was characterized by higher cadence and shorter (average) step and stride lengths compared to overground walking. This agrees with many previously published studies (Alton et al., 1998; Dingwell et al., 2001; Riley et al., 2007). As overground walking is the ecologically valid mode of human walking, increases in speed are achieved through coordinated increases in both step/stride lengths and cadence, whereas treadmill walking constrains step/stride length modulation and relies more heavily on increasing cadence.

Considering our explorative questions, we saw significant differences in basic anthropometrics between the males and females. The main effect of sex on the gait parameters is consistent with established anthropometric differences between them. As the females had a shorter stature and lower limb lengths, they adopted shorter steps and faster cadence to achieve a given walking speed (Ko et al., 2011; Abualait & Ahsan, 2022; Rayner et al., 2024). Importantly, these differences persisted even when speeds were matched.

If we look at some previously published literature, Parvataneni et al., (2009) and Berryman et al., (2012) reported higher metabolic cost on treadmill compared to overground in healthy OA. In this study by focusing exclusively on healthy YA, we show that the metabolic penalty is not exclusive to aging and rather may arise from more fundamental demands of treadmill walking. In previous work (Das Gupta et al., 2021) we did not see an elevation in NCoW for our YA at PWS. Our current results suggest that this apparent equivalence may not generalize beyond PWS and so in this study we saw an elevated metabolic cost for all the three tested walking speeds, despite the CWS of 1.3 m/s being very close to the general PWS of humans. A recent systematic review and meta-analysis also reported increased relative oxygen consumption and cadence, and reduced stride/step lengths on treadmill compared to overground walking in healthy adults (Vickery-Howe et al., 2023), thus showing similarity with our results. The spatiotemporal shift in gait suggests that participants show a cautious strategy and may have prioritized stability over energetic economy. Such strategies likely increase muscle activation and control effort, analogous to what is seen in conditions which are novel or where balance is challenged (Donelan, Kram & Kuo, 2001). While we did not record Electro-Myo-Graphy (EMG), prior work in OA (Peterson & Martin, 2010) indicates that treadmill walking increases antagonistic muscle co-activation, which can be seen as a compensatory strategy to achieve lower limb joint stability, often at the cost of a metabolic penalty. Even in healthy YA, like in this study, more stabilizing muscle activity may have driven up the metabolic cost.

Another interesting finding of this study is the divergent behavior of GCoW and NCoW. GCoW was elevated on the treadmill but did not vary significantly with speed and exhibited the classic U-shaped relationship; whereas NCoW increased with speed and was also elevated on the treadmill. Since GCoW represents the whole-body level energy required to travel a unit distance and has repeatedly been shown to exhibit a minimum near PWS/CWS, it reflects the ecological and energy minimization theories (Ralston, 1958; Zarrugh et al., 1974; Minetti & Alexander, 1997) that govern human walking. In contrast, for NCoW, it is expected that we do not see the U-shaped relationship between cost and speed. This is because we measure RMR only at the beginning and subtract the same RMR value from the GEE at every walking speed to calculate the NEE and then divide it by the different walking speeds to get the NCoW. At slow speeds, the RMR is larger and at faster speeds it is smaller, relative to walking energy expenditure. As such when we subtract the same RMR from the GEE, at slower speeds, the NEE is disproportionately reduced, thereby suppressing the NCoW. At faster speeds the similar subtraction does not impact the NEE substantially and so the NCoW increases.

Our study has several limitations. Firstly, we only measured healthy Japanese YA and so we cannot generalize to older, clinical, or less fit (patient) populations and possibly to other ethnicities. Secondly, though we gave each one of our participants at least ten minutes of treadmill familiarization in the beginning, we only did it at the CWS and this might not eliminate all novelty and adaptation effects. In Das Gupta et al., (2021) we gave similar amount of treadmill familiarization at PWS and saw that it was enough for the healthy YA in that study. So, we can possibly claim that familiarization was not the main driver for the increased MCoW for YA on treadmill in this study. Finally, overground walking was paced using visual and auditory cues and participants had to take turns, which may reduce natural gait variability and slightly alter gait parameters. However, we advised the participants to take smooth turns and visually ensured that they were doing so. All the participants were also walking at the desired speed and were in their metabolic steady state after 2 minutes of overground walking. Altogether, we think that these limitations do not substantially affect and change our primary conclusions about YA and the consistent effect of the treadmill compared to overground is much more dominant.

Looking forward, future studies should aim to combine metabolic measurement with EMG and kinetics and kinematics from motion capture to identify muscle activation patterns, joint work, postural stability metrics and their (relative) contributions to increased metabolic cost. This may help establish causal links between the mechanics and energetics of human walking. Investigating short term (days) and long term (weeks to months) adaptations to treadmill walking and assessing whether repeated exposure reduces the energetic penalty on treadmills is also fruitful to study. Studies manipulating sensory feedback, such as visual flow via virtual reality, proprioceptive inputs via musculo-tendon vibrations and vestibular control via Electronic Vestibular Stimulation would furthermore be interesting to study how sensory context and possible reweighting on treadmills compared to overground modifies gait and thereby lead to the metabolic penalty.

## Conclusion

To the best of our knowledge, this study is one of the few recent investigations that provides strong evidence that even healthy young adults incur a substantial and consistent energetic penalty on treadmills compared to overground. They also exhibit an altered and possibly sub-optimal gait when walking on a treadmill, across a range of matched speeds. Based on these findings, we caution both researchers and practitioners against the uncritical use of treadmill data to infer overground energetic and gait behavior as it may not be a proxy for the ecologically valid overground walking. This is critically important for advancing scientific understanding, informing evidence-driven recommendations for future clinical and rehabilitation applications and for conducting improved modeling studies trying to connect the mechanics and energetics of human walking, using empirical data.

## Acknowledgements

This is an independent research funded by the internal grant of the National Institute of Advanced Industrial Science and Technology (AIST), Japan and conducted at the same institute. Ms. Kanako Kamishita was supported by a Shoshisha Foundation’s scholarship. Additionally, Dr. Sauvik Das Gupta was supported by a JSPS KAKENHI, Grant-in-aid for Early-Career Scientists (ECS) award, (Project number: 25K20863) while this study was being conducted. The authors would like to thank Mrs. Saori Ohtani for her help during the data collection sessions.

## Author contributions

SDG and KK performed the experiments. SDG did the data analysis and made the table and figures. SDG conceptualized and designed the study, drafted the manuscript and revised it. SDG, KK, MK and YK were involved in the manuscript development and revision. YK had the internal funding to perform the study.

## Availability of data and materials

The datasets generated during and/or analyzed during the current study are generally protected (due to AIST institutional guidelines and ethical approval) and are not openly available. However, they maybe available from the last author (YK) on reasonable request.

## Code availability

Not applicable.

## Declarations

### Conflict of interest

The authors declare no competing interests.

## Supplementary Figure

**Figure 1:**
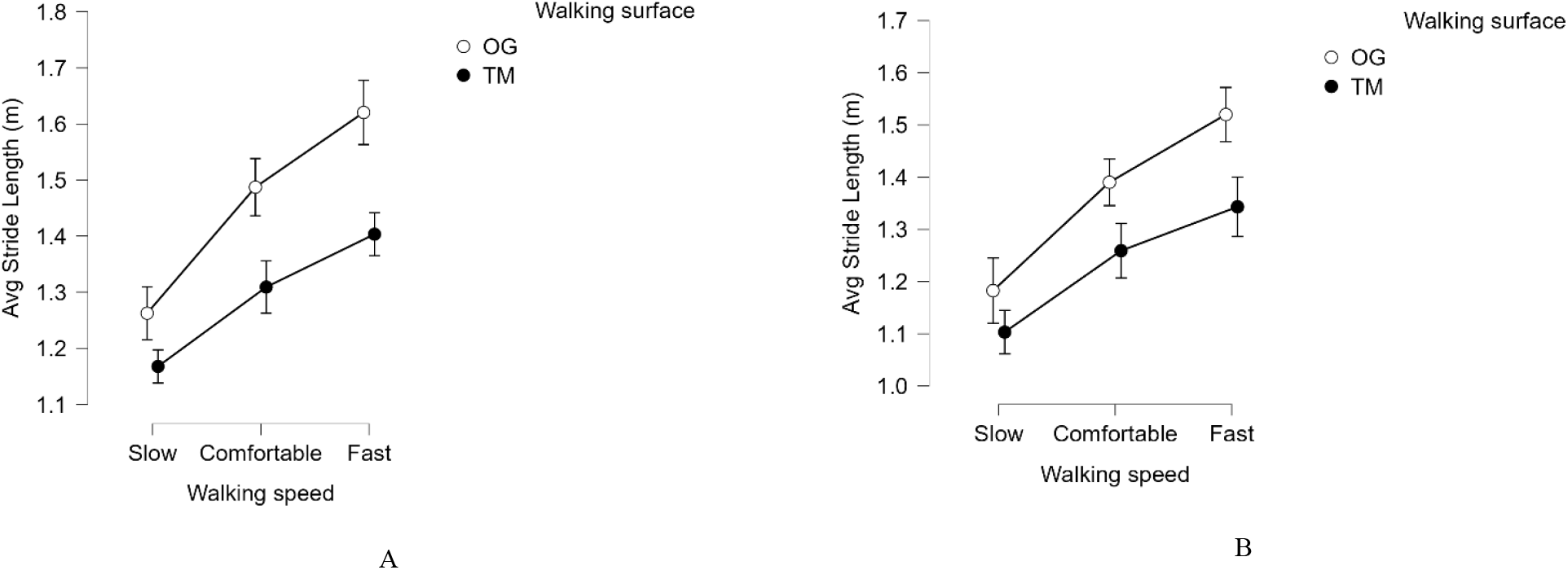
Line-plots of average stride length are shown for overground (OG, in open white circles) and treadmill (TM, in black closed circles). The circles denote the mean values and the vertical straight black lines across the circles are the error bars denoting the 95% CIs. The x-axis shows the walking speeds in text and the y-axis shows the gait parameter and its unit. Figure 1A depicts the data for the young males and Figure 1B depicts the data for the young females. It is clear from these figures that at matched speeds, the treadmill forces a reduced average stride length (smaller strides) for both males and females and with increasing speeds, expectedly there is an increase in average stride length. If we compare the y-axis values of figures 1A and B, we can see that the females had a shorter average stride length than males.

## Notes

### Competing Interest Statement

The authors have declared no competing interest.

